# Comprehensive Network and Structural Analysis of Bovine Papillomavirus, Squamous Cell Carcinoma Markers, and Elucidation of Efficacy Mechanisms of Phytochemicals from *Thuja Occidentalis*

**DOI:** 10.1101/2023.12.05.570096

**Authors:** Shafiqur Rahman, Arun HS Kumar

## Abstract

Papillomaviruses infect cutaneous tissue in various species including bovines and from benign warts to malignant squamous cell carcinoma causing severe economic losses to the farmers. The mechanisms by which bovine papillomaviruses interact with host tissue are unclear. Hence in this study using classical network analysis tools, we evaluated interactions of Bovine papilloma (BPV) variants, with markers and receptors implicated in squamous cell carcinoma. Additionally, the thuja phytoconstituents were also evaluated for its potential to target the BPV and squamous cell carcinoma network interactions to understand the mechanism of its clinical benefits. Various protein composition of 14 different virus variants of BPV were assessed against 24 markers of squamous cell carcinoma. Among these interactions EGFR consistently exhibited high-affinity interactions with the E1 protein in all isoforms of BPV. Type 4 BPV displayed the maximum number of binding sites (14) with a binding pocket score ranging from 15.47 to 141.34 and a probability score of 0.75 to 0.99. The comparison of the binding pockets identified that BPV types 2 and 13 had the highest number of common amino acid sequences. Further the alpha helix structure of specific common amino acid sequences, contribute to a more robust and widespread affinity interaction with both E1 of various BPV types and EGFR. Analysis of thuja phytochemicals suggested superior efficacy of Beyerene and Terpinene-4-ol towards all ten BPV targets and bEGFR. In conclusion, our comprehensive study leading to identification of E1 protein of BPV as a major interacting network with bEGFR, their key binding sites, and efficacy of thuja phytoconstituents offer valuable insight into further experimental validation and development of novel therapeutic strategies against BPV-associated diseases.

## Introduction

Papillomaviruses are a group of oncogenic DNA tumor viruses that can infect cutaneous and mucous epithelia in various species including bovines.^[1-4]^ Upon infection, they typically induce the formation of papilloma’s or warts, which generally regress without causing significant clinical problems in the host. The intriguing aspect of papillomavirus biology lies in its close association with the host epithelial cell’s differentiation process.^[1-7]^ Despite the predominantly benign nature of papillomavirus infections, there are instances where benign warts persist and may progress to squamous cell carcinoma.^[5, 7-10]^ This progression to malignancy although is reported to be a rare outcome, represents a dead-end process for the virus and host causing severe economic losses to the farmers.^[1, 7, 11]^ Neoplastic progression involves the transformation of infected cells into cancerous cells, and it is a complex phenomenon influenced by both viral and host factors. Understanding the molecular events governing the papillomavirus and neoplastic progression is crucial for developing strategies to prevent and treat associated diseases, including squamous cell carcinoma.

Bovine Papillomavirus (BPV) infections in cattle have garnered significant attention due to their association with squamous cell carcinoma and other pathological conditions.^[1, 5, 6, 12]^ Understanding the diverse array of BPV variants, their interactions with host cellular components, and exploring potential therapeutic interventions is crucial for advancing our knowledge in veterinary medicine and developing effective strategies to manage these viral infections. In veterinary medicine, certain papillomavirus types have gained prominence for their strong association with malignant neoplasms. BPV types 1, 2, 4, and 13, as well as feline papillomavirus (FcaPV) types 2 and 3, have recently been identified as significantly correlated with the development of malignant conditions such as squamous cell carcinoma (SCC), bowenoid in situ carcinoma (BISC), and transitional cell carcinoma.^[5, 11, 13]^ While the link between these papillomavirus types and malignant neoplasms is established, the precise role of bovine papillomaviruses in the development of precursor lesions and the subsequent progression of these lesions to SCCs remains a subject of debate within the veterinary community.^[10, 11]^ The specific causative agent responsible for this progression has not been conclusively identified to date. Research in this area is ongoing, aiming to elucidate the intricate mechanisms underlying the relationship between these papillomaviruses and the development of malignant lesions in animals. Understanding the nuanced interplay between viral factors and host biology is crucial for unraveling the complexities of papillomavirus-associated carcinogenesis. In this context, using classical network analysis tools, the present study integrates evaluations of BPV variants, their interactions with markers and receptors implicated in squamous cell carcinoma to identify the major mechanisms of associations between BPV and squamous cell carcinoma. Additionally, the thuja phytoconstituents which have shown considerable clinical efficacy were also evaluated for its potential to target the BPV and squamous cell carcinoma network interactions to understand the mechanism of clinical benefits.

## Materials and Methods

All the reported proteins in different types of Bovine papillomaviruses (BPV) were identified and reviewed in the UniProt database and their 3D structure if not already available were generated by homology modelling. The targets for which 3D structure was not available, the protein sequence (FASTA format) was extracted from the UniProt database, and the 3D structure were constructed by homology modelling using the SWISS-MODEL server (https://swissmodel.expasy.org/). For generating the 3D structures in the SWISS-MODEL server, each of the protein sequence in FASTA format was loaded into the SWISS-MODEL server and the molecular modelling was initiated to generate PDB format of the protein. Different markers/receptor assisting BPV to establish and cause squamous cell carcinoma in bovines (bSCC) reported in the literature were identified and considered for analysis in this study. The PDB code, AlphaFold or homology modelled structure of the all the BPV proteins and their reported targets in bSCC was used to import the protein structure onto the Chimera software and the number of hydrogen bonds (H-bond) formed between BPV proteins and their targets in bSCC at 10 Armstrong (10A) distance was evaluated. A heatmap of the number of H-bonds formed between BPV proteins and their targets was generated to identify the high affinity interactions. ^[14-16]^

E1 and EGFR were observed to show high affinity interactions in all isoforms of BVP. Hence, the binding sites of E1 of various BPV and EGFR were identified using the PrankWeb: Ligand Binding Site Prediction tool (https://prankweb.cz/). Analysis of binding sites sequence of various types of BPV were done to identify the key amino acid sequence and subsequently their common amino acid sequence were compared amongst various types of BPV. Briefly, the sequence of the top binding site (based on score and probability value) were selected for further analysis. The top binding site of E1 of BPV type 1 was compared with all other BPV variants.

A peptide sequence having at least 7 common amino acids was taken into consideration for further analysis. The 3D structure (PDB format) of these peptide sequences were prepared in Chimera software and then these structures (both alpha helix and antiparallel structures were generated) were tested for their interaction with E1 of all BPV and EGFR using similar criteria reported above. A heatmap of the number of H-bonds formed between different peptides and its associated network E1 proteins of various types of BPV was generated to identify the high affinity interactions.

The phytoconstituents of Thuja which is reported to have clinical efficacy in the treatment of BVP, were evaluated for the affinity against E1 of various BPV and EGFR by molecular docking using AutoDock vina 1.2.0 as reported before.^[14-17]^

## Results

In the UniProt database, an analysis of reported proteins from various types of Bovine Papillomavirus (BPV) revealed the existence of 14 different virus variants (Figure 1a). These distinct BPV types exhibited varying protein compositions, each demonstrating an affinity for binding to different markers or receptors. Notably, type 4 displayed the highest number of proteins (11), followed by type 2 (10), type 1, 6, and 13 (8 each), type 3, 5, 28, and 29 (7 each), and type 10 (2). Conversely, type 7, 8, 9, and 12 each exhibited only a single protein (Figure 1b). In this study, we conducted an analysis of 24 markers/receptors associated with BPV-induced squamous cell carcinoma in bovines. These markers included VEGF-C, EGFR, Ki67, PCNA, MMP-2, 7, 9, 14, TP53, Caspase-3, AIF, ATG-5,9, PDGFC, PDGFC-alpha, TIMP-1,2, KRT-5, 13, CCL2, GPC1, bFGF, IVL, and FLG (Figure 1a). Our interaction study revealed a higher affinity for VEGF-C, EGFR, PDGFC, PDGFC-alpha, bFGF, GPC1, and FLG with the majority of BPV variants (Figure 1b). Notably, BPV types 8, 10, and 12 showed no significant interaction with the markers, while most proteins from other types exhibited binding affinity with multiple markers. BPV types 7 and 9 demonstrated an affinity for all the markers/receptors. Among the markers/receptors, EGFR consistently exhibited high-affinity interactions with the E1 protein in all isoforms of BPV. Consequently, we identified the binding sites of E1 in various BPV types with EGFR. Type 4 displayed the maximum number of binding sites (14) with a binding pocket score ranging from 15.47 to 141.34 and a probability score of 0.75 to 0.99. In comparison, type 6 had 12 binding sites with lower binding and probability scores (7.77-83.98 and 0.45-0.99, respectively) than types 28 and 29, which shared similar binding sites with higher scores. Types 1, 2, 3, 5, and 13 had an equal number of binding sites (11), but type 5 exhibited a higher binding pocket score (15.47-141.34) and probability score (0.75-0.99) than the others. EGFR itself demonstrated only four binding sites with a binding pocket score ranging from 1.06 to 26.78 and a probability score from 0.00 to 0.89 (Figure 2).

**Figure 1.**
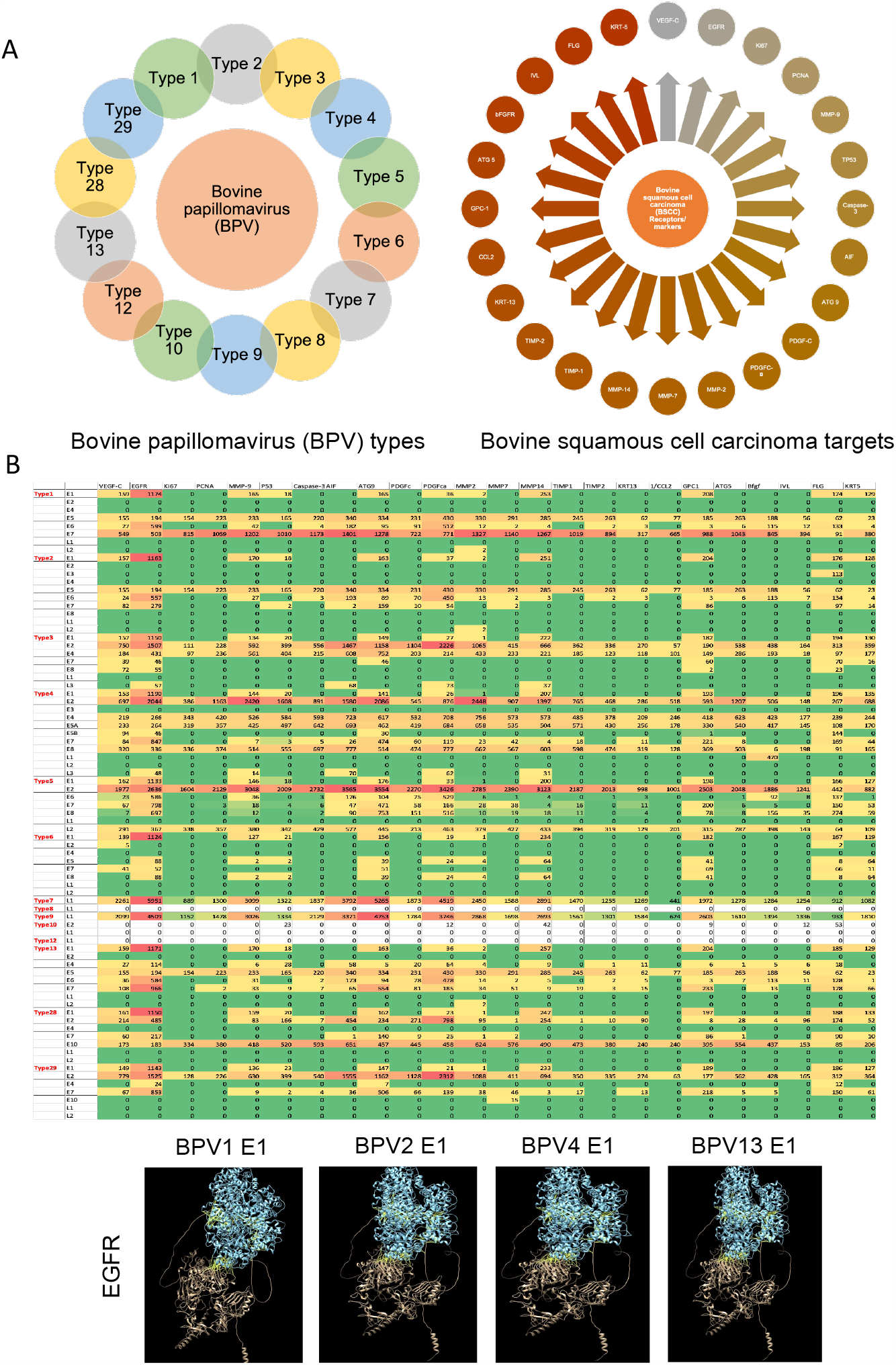
A) Various isotypes of bovine papilloma virus (BPV) used in this study analysis and various receptors/markers associated with bovine squamous cell carcinoma (bSCC). B) Heat map showing interactions (hydrogen bonds) between surface markers of BPV isotypes and bSCC. Panel below shows representative images of E1 protein of various BPV variants interacting with bEGFR.

**Figure 2.**
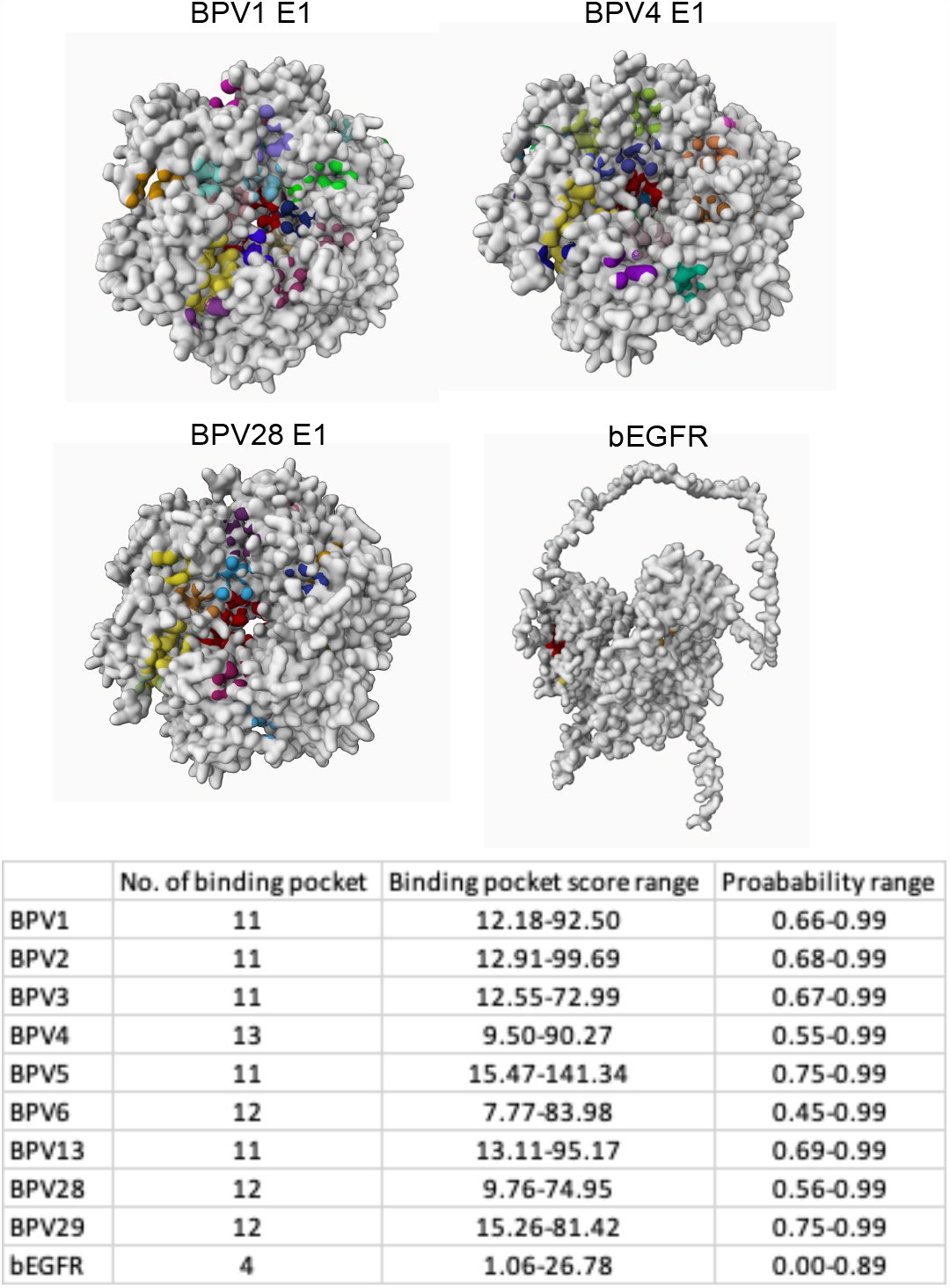
Binding pockets of E1 protein of various BPV variants (type 1, 4 and 28) and bEGFR. The total number of binding pockets of all BPV variants, the range of their binding pocket and probability scores are summarized in the table below.

To elucidate the key amino acid sequences in the major binding sites of various BPV types, selected based on their score and probability values (Figure 2), a comparative analysis was undertaken. Subsequently, we aimed to identify common amino acid sequences among different types of BPV. The top binding site of E1 in BPV type 1 was compared with the corresponding sites in all other BPV variants. BPV types 2 and 13 exhibited the highest number of common amino acid sequences, with four identified sequences. Following closely were types 4, 6, and 28, each sharing two common amino acid sequences. Conversely, types 5 and 29 did not demonstrate any common amino acid sequences with the E1 of BPV type 1 (Figure 3). This comparative analysis provides insights into the shared molecular features among different BPV types, shedding light on potential conserved elements in their binding sites.

**Figure 3.**
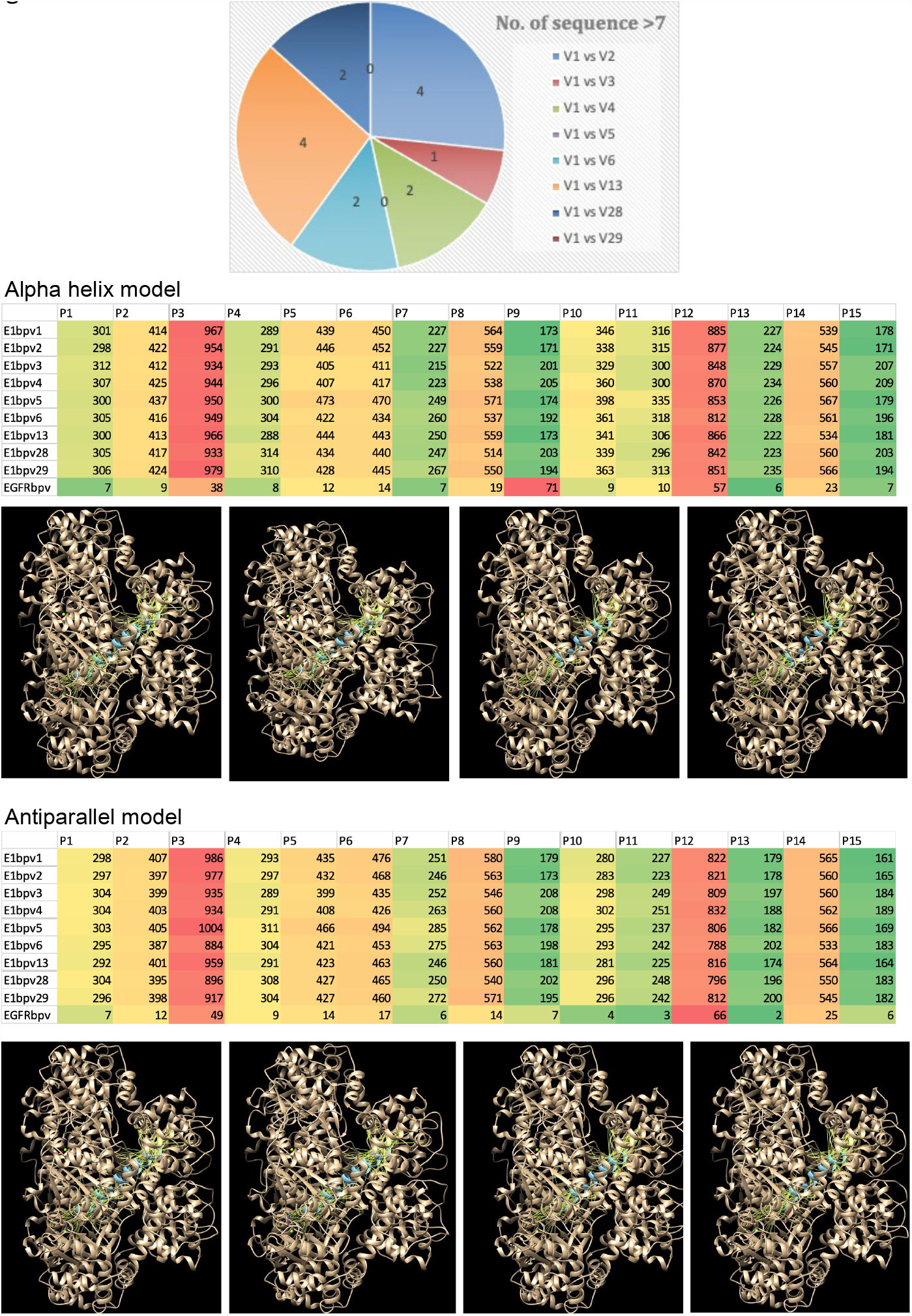
The similarities in the sequences of the binding pockets of E1 protein of BPV type 1 with all other types. The number of similar sequences between each comparison are shown in the wheel diagram. The structures of the similar sequences were generated as either alpha-helix or antiparallel model and checked for interactions (Hydrogen bonds) with all BPV variants and bEGFR. The heatmaps of these interactions are presented. A few representative images of these interactions are also shown.

The interaction analysis of common amino acid sequences with both E1 of all BPV types and EGFR was conducted utilizing both alpha helix and antiparallel structures. Specifically, the alpha helix structure formed by the common amino acid sequence for type 1 and 13 BPV, particularly the 3rd sequences, exhibited the highest affinity interaction with all other BPV variants (Figure 3). This was followed by the 2nd sequence of type 6 and 28 BPV. Notably, the 2nd sequences of type 6 and 28 BPV showed the least affinity interaction in comparison. This observed pattern of affinity interaction was consistent for the antiparallel structure as well. The findings suggest that specific amino acid sequences, particularly those within the alpha helix structure, contribute to a more robust and widespread affinity interaction with both E1 of various BPV types and EGFR. Further exploration of these structural nuances may provide valuable insights into the molecular mechanisms underlying these interactions.

In our study, we aimed to assess the potential efficacy of thuja against BPV by evaluating its major phytoconstituents against key BPV targets and EGFR. We identified six major phytoconstituents—alpha-Thujone, beta-Thujone, Fenchone, p-Cymene-8-ol, Terpinene-4-ol, and Beyerene—from the literature. The relative concentrations of these phytoconstituents are summarized in Figure 4, with alpha-Thujone being the predominant constituent, constituting approximately 70% of the thuja extract. The remaining phytoconstituents collectively make up less than 10% of the extract. Conducting molecular docking studies of these phytoconstituents against the screened targets revealed significant affinity of Beyerene and Terpinene-4-ol towards all ten targets tested, including bEGFR (Figure 4). Notably, alpha and beta-Thujone exhibited substantial affinity against BVP type 4/5 E1 and bEGFR. Fenchone displayed major affinity exclusively against bEGFR, while p-Cymene-8-ol showed the least affinity against all ten targets examined (Figure 4). Of all the phytoconstituents in thuja, Beyerene emerged as the most promising, displaying the highest affinity against all ten targets tested (Figure 4). These findings suggest that thuja and its constituents, particularly Beyerene, may hold potential for therapeutic intervention against BPV and EGFR-related conditions. Further exploration and validation of these molecular interactions could provide valuable insights into the development of novel therapeutic strategies.

**Figure 4.**
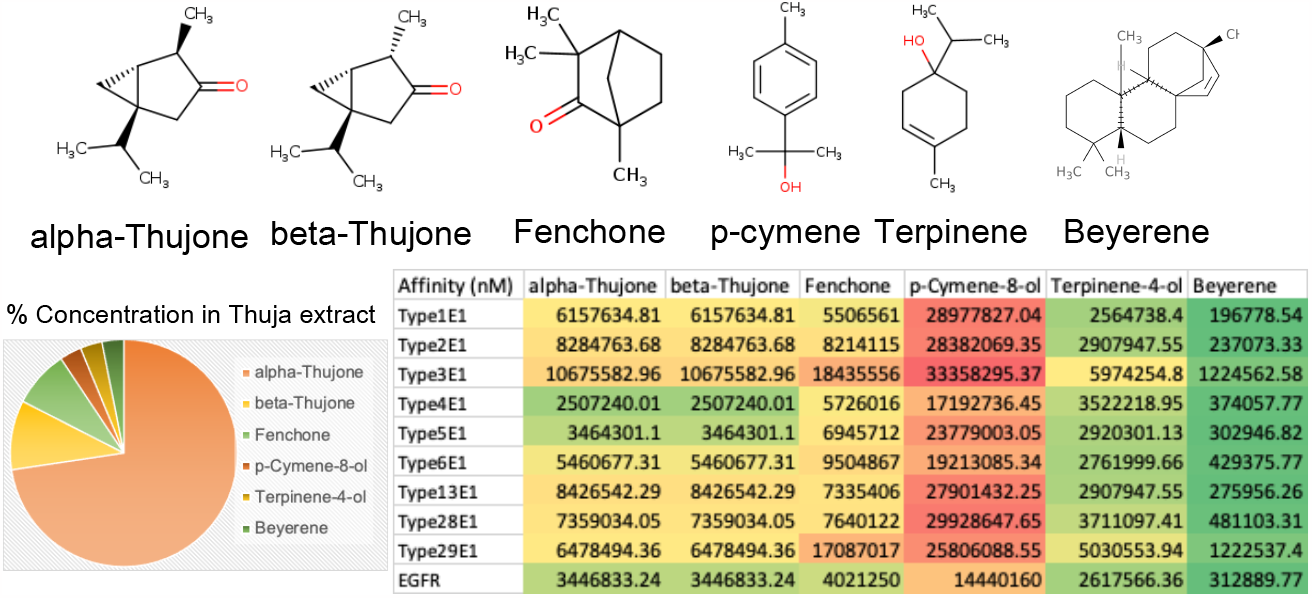
The structure of thuja phytoconstituents are presented. The relative composition of each of the phytoconstituents in a thuja extract is shown in the wheel diagram. The affinity of the thuja phytoconstituents against E1 protein of various BPV variants and bEGFR is presented as heatmap.

## Discussion

This study expands beyond the viral focus to scrutinize host cellular components associated with BPV-induced squamous cell carcinoma.^[13, 18-22]^ A panel of 24 markers and receptors, including crucial entities such as VEGF-C, EGFR, and TP53, was systematically analyzed to discern their interactions with various BPV variants. The outcomes of this analysis shed light on potential key players in the viral-host interplay, which can help development of targeted therapeutic interventions. Furthermore, this study delves into the intimate interaction between the E1 protein of BPV and EGFR,^[20, 23, 24]^ a central player in cellular signaling pathways. Identification of binding sites and the comparative analysis of common amino acid sequences across different BPV types unravel potential conserved elements, offering a deeper understanding of the molecular basis of these interactions. Extending the exploration of natural therapeutics, the study also investigated the major phytoconstituents of thuja for their potential efficacy against BPV-EGFR networks due to their prior reports of clinical efficacy.^[25-28]^ The focus on alpha-Thujone, beta-Thujone, Fenchone, p-Cymene-8-ol, Terpinene-4-ol, and Beyerene provides a nuanced view of the phytochemical composition of thuja. Molecular docking studies against key BPV targets and EGFR elucidate the potential of these phytoconstituents in mitigating the pathogenesis of the virus. The comprehensive analysis of BPV variants and their interactions with key markers, receptors, and phytoconstituents from thuja provides valuable insights into potential therapeutic avenues and molecular mechanisms. We delve into the implications of our findings and their relevance to the understanding of BPV-induced squamous cell carcinoma in bovines. Firstly, the diversity in the protein compositions of different BPV types, as identified through the UniProt database, highlights the intricate nature of viral variants. The varying numbers of proteins and their affinities for different markers/receptors underscore the complex interactions between BPV and host cellular components. Notably, the lack of significant interactions for BPV types 8, 10, and 12 with the screened markers suggests distinct characteristics and potential variations in their pathogenic mechanisms.^[5, 6, 18]^

The analysis of 24 markers/receptors associated with BPV-induced squamous cell carcinoma provides a comprehensive overview of potential molecular targets. The higher affinity interactions observed for VEGF-C, EGFR, PDGFC, PDGFC-alpha, bFGF, GPC1, and FLG across multiple BPV variants indicate the significance of these markers in the context of viral pathogenesis.^[5, 6, 18]^ Noteworthy is the consistent high-affinity interaction of EGFR with the E1 protein, emphasizing its central role in BPV-related molecular pathways. The identification of binding sites in E1 proteins for various BPV types and their comparison reveals potential conserved elements and distinct structural features. The differential binding characteristics, particularly the maximum number of binding sites in type 4, offer insights into the diversity of viral-host interactions. Such information can guide further studies on the functional significance of these binding sites in the context of viral replication and pathogenesis.^[1, 2, 29]^ The comparative analysis of common amino acid sequences in the major binding sites of various BPV types further elucidates shared molecular features. The identification of common sequences, especially in types 2 and 13, suggests potential conserved regions crucial for interactions with host cellular components. Understanding these commonalities could pave the way for the development of targeted therapies against multiple BPV variants. The structural analysis of common amino acid sequences using both alpha helix and antiparallel structures provides additional depth to our understanding of the molecular interactions. The observed patterns of affinity interactions highlight the importance of specific amino acid sequences, especially those within the alpha helix structure, in mediating robust interactions with both E1 of various BPV types and EGFR.

The evaluation of thuja phytoconstituents against BPV targets and EGFR, the predominance of alpha-Thujone in the extract underscores its potential as a major contributor to the observed effects.^[25-28]^ The molecular docking studies reveal promising affinities of Beyerene and Terpinene-4-ol against all tested targets, suggesting their potential as key therapeutic components. The distinct affinities of alpha and beta-Thujone against specific BPV types and EGFR, along with the unique characteristics of Fenchone and p-Cymene-8-ol, provide a basis for further investigation into the therapeutic potential of thuja phytoconstituents against BPV-induced conditions.

In conclusion, our comprehensive study contributes to the understanding of BPV diversity, molecular interactions, and the potential therapeutic efficacy of thuja phytoconstituents. This multidimensional approach contributes to the ongoing efforts to understand BPV-associated diseases and underscores the importance of translational research in veterinary medicine. The identified markers, binding sites, and phytoconstituents offer valuable targets for further experimental validation and development of novel therapeutic strategies against BPV-associated diseases.

## Acknowledgements

Research support from University College Dublin-Seed funding/Output Based Research Support Scheme (R19862, 2019), Royal Society-UK (IES\R2\181067, 2018), Stemcology (STGY2917, 2022), SKUAST-Jammu, and NAHEP-IDP (ICAR New Delhi), is acknowledged.

